# Programming large target genomic deletion and concurrent insertion via a prime editing-based method: PEDAR

**DOI:** 10.1101/2021.05.12.443800

**Authors:** Tingting Jiang, Xiao-Ou Zhang, Zhiping Weng, Wen Xue

## Abstract

Genomic insertions, duplications, and insertion/deletions (indels) account for ~14% of human pathogenic mutations. Current gene editing methods cannot accurately or efficiently correct these abnormal genomic rearrangements, especially larger alterations (>100 bp). Thus, developing a method to accurately delete insertions/duplications and repair the deletion junction could improve the scope of gene therapies. Here, we engineer a novel gene editor, PE-Cas9, by conjugating Cas9 nuclease to reverse transcriptase. Combined with two prime editing guide RNAs (pegRNAs) targeting complementary DNA strands, PE-Cas9 can direct the replacement of a genomic fragment, ranging from to ~1-kb to >10-kb, with a desired sequence at the target site without requiring an exogenous DNA template. In a reporter cell line, this <u>PE</u>-Cas9-based <u>d</u>eletion <u>a</u>nd <u>r</u>epair (PEDAR) method restored mCherry expression through in-frame deletion of a disrupted GFP sequence. We further show that PEDAR efficiency could be enhanced by using pegRNAs with high cleavage activity or increasing transfection efficiency. In tyrosinemia mice, PEDAR removed a 1.38-kb pathogenic insertion within the *Fah* gene and precisely repaired the deletion junction to restore FAH expression in liver. This study highlights PEDAR as a tool for correcting pathogenic mutations.

## Main

Genetic insertions, duplications, and indels (<u>in</u>sertion/<u>del</u>etion) account for ~14% of 60,008 known human pathogenic variants^1^ (**Fig. 1a**). Many of these abnormal insertions and duplications involve larger DNA fragments (>100 bp). Indeed, retrotransposon element insertions, ranging from 163 to 6000 bp^2, 3^, disrupt the normal expression and function of genes^4^ thereby causing genetic diseases like cystic fibrosis, hemophilia A, X-linked dystonia-parkinsonism, and inherited cancers^4–7^. Precise genome editing technologies that simultaneously delete the inserted or duplicated DNA sequences and repair the disrupted genomic site might provide a way to treat a wide range of diseases.

**Fig 1.**
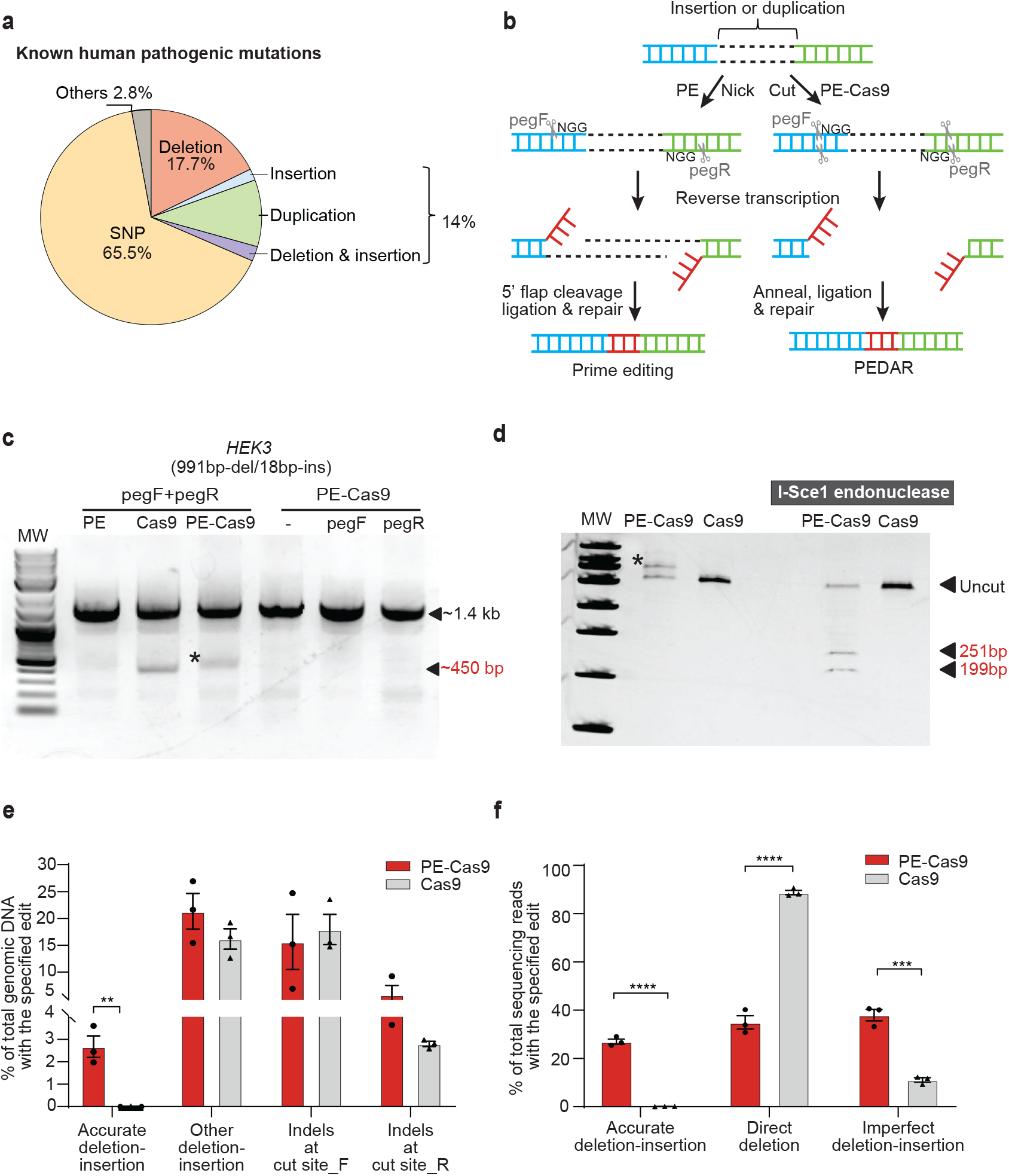
PEDAR mediates large target deletion and simultaneous insertion at an endogenous genomic locus. **a,** Classification of the 60,008 known human pathogenic genetic variants reported in the ClinVar database^1^. **b,** Overview of using prime editing (left) and PEDAR (right) to generate accurate deletion-insertion. Prime Editing: Two PE complexes – consisting of pegRNA (pegF or pegR) and a Cas9 nickase (Cas9 H840A) conjugated to an engineered reverse transcriptase (RT) – recognize ‘NGG’ PAM sequences, bind, and nick the target DNA strands. After hybridization of the nicked DNA strand to the primer binding sequence of pegRNA, the desired edit is reverse transcribed into the target site using the RT template at the 3’ extension of pegRNA. The desired inserted edits (red) at the two nicking sites are complementary. After equilibration between the edited 3’ flap and the unedited 5’ flap, the 5’ flap is cleaved, and DNA repair results in coupled deletion and insertion of target sequence. PEDAR: Dual PE-Cas9:pegRNA (pegF or pegR) complexes recognize ‘NGG’ PAM sequences, bind, and cut the target DNA. The two complementary desired edits (red) are reverse transcribed into the target sites using the RT template at the 3’ extension of pegRNAs. The inserted sequences are annealed, and the doublestranded DNA break is repaired **c,** Deleting a 991-bp DNA fragment and simultaneous insertion of I-Sce1 recognition sequence (18bp) at the *HEK3* locus (Chr9:107422166-107423588). Target genomic region was amplified using primers that span the cut sites. The paired pegRNAs targeting complementary DNA strand are denoted as pegF and pegR. HEK293T cells were transfected with PE, Cas9, or PE-Cas9 with or without single or paired pegRNAs. The ~450-bp band is the expected deletion amplicon (denoted with *), and the ~1.4-kb band is the amplicon without deletion. **d,** Deletion amplicons from Cas9- or PE-Cas9-treated groups shown in (c) were incubated with or without I-SceI endonuclease and analyzed in 4-20% TBE gel. Digested products are marked by arrows with expected sizes. Original amplicon is marked as “uncut”. The band with insertion of i-Sce1 recognition sequence is denoted with *. **e,** PE-Cas9 or Cas9-mediated absolute rates of all the editing events in total genomic DNA at *HEK3* site. Data represent mean± SEM (n = 3 biologically independent samples). P= 0.0053 (**). **f,** Deep sequencing of deletion amplicons shown in (c). Bar chart shows distribution of all deletion events, including accurate deletion-insertion, direct deletion (deletion without any insertions), or imperfect deletion-insertion. Editing rate = the reads with indicated editing / total deletion events. Data represent mean ± SEM (n = 3 biologically independent samples). P=<0.0001 (****), 0.002 (***), two-tailed t-test.

The CRISPR/Cas9 system is a powerful gene editing tool for correcting pervasive pathogenic gene mutations. Using dual single guide RNAs (sgRNA), Cas9 can induce two double-strand breaks (DSBs). The two cut ends are then ligated through the non-homologous end joining (NHEJ) repair pathway, leading to ≤5-Mb target fragment deletion in vitro^8, 9^ and in vivo^10–12^. However, the random indels generated by NHEJ lower the editing accuracy of this method. When a donor DNA template is present, CRISPR/Cas9 can insert a desired sequence at the cut site to more accurately repair the deletion junction through homology directed repair (HDR)^13^. This method has been used successfully in precise gene deletion and replacement applications^14^. Nevertheless, the repair efficiency of CRISPR-mediated HDR is hindered by the exogenous DNA donor and is limited in post-mitotic cells^15, 16^. To further expand the gene editing toolbox, a novel CRISPR-associated gene editor – called prime editing (PE)^17^ – was developed by conjugating an engineered reverse transcriptase (RT) to a catalytically-impaired Cas9 ‘nickase’ (Cas9^H840A^) that cleaves only one DNA strand. An extension at the 3’ end of the <u>p</u>rime <u>e</u>diting <u>g</u>uide RNA (pegRNA) functions as an RT template, allowing the nicked site to be precisely repaired^17, 18^. Thus, PE can mediate small deletion, insertion, and base editing without creating DSBs or requiring donor DNA^17^, and holds great promise for correcting human genetic diseases^19–22^. Yet, PE has not been applied to delete larger DNA fragments.

To achieve accurate and efficient large fragment deletion and simultaneous insertion without requiring a DNA template (**Supplementary Fig. 1a**), we modified the prime editing system to employ a pair of pegRNAs (hereafter referred to as pegF and pegR) rather than one pegRNA and one nicking guide RNA. We reasoned that using two pegRNAs would enable concurrent targeting of both DNA strands. At the 3’ extension of each pegRNA is a reverse-complementary RT template, which encodes the sequences for desired insertion. In theory, this newly-engineered system could mediate accurate deletion-repair through the following steps (**Fig. 1b**, left side): (i) prime editor recognizes the ‘NGG’ PAM sequence, binds, and nicks the two complementary strands of DNA on either side of the large fragment^8^; (ii) the desired insertion sequences are reverse transcribed into the target site using the RT template linked to the pegRNAs; (iii) the complementary DNA strands containing the edits are annealed; (iv) the original DNA strands (i.e., 5’ flaps) are excised; and (v) DNA is repaired. However, Cas9 nickase cannot not effectively mediate larger target deletions with paired guide RNAs^23, 24^. Indeed, PE applications reported in the literature are limited to programing deletions of less than 100 bp, raising the concern that PE cannot generate long genomic deletions^18^.

Fully active Cas9 nuclease has been used to program larger deletions with dual sgRNAs^14^. Therefore, we conjugated an active Cas9 nuclease, instead of Cas9 nickase, to the RT^17^ to create “PE-Cas9” (**Supplementary Fig. 1b**). With a single pegRNA^17^, PE-Cas9 and PE generated similar rates of 3-bp CTT insertion at the cut/nicking site of an endogenous locus (**Supplementary Fig. 1c**), indicating that Cas9 nuclease activity does not affect prime editing efficiency. We hypothesized that, with the guidance of two pegRNAs targeting both complementary strands of DNA, PE-Cas9 can introduce two DSBs and delete the intervening DNA fragment between the two cut sites. Concurrently, the desired edits are incorporated at target sites using the RT template at the 3’ extension of the pegRNAs. The two complementary edits then function as homologous sequences to direct the ligation and repair of the deletion junction. We term this method “PE-Cas9-based deletion and repair” or PEDAR (**Fig. 1b**, right side).

We compared the efficiency of PEDAR, PE, and Cas9 systems in coupling large target deletion and accurate insertion at the endogenous *HEK3* genomic locus in HEK293T cells. We designed two pegRNAs with an offset of 979 bp (distance between the two ‘NGG’ PAM sequences) to program a 991bp-deletion/18bp-insertion at the *HEK3* site. The RT template at the 3’ extension of the pegRNAs encodes an I-SceI recognition sequence (18-bp), which will be reversed transcribed and integrated into the target site (**Supplementary Fig.1d**). Paired pegRNAs along with PE, PE-Cas9, or Cas9 were transfected into cells. Delivery of PE-Cas9 with or without single pegRNA was used as a negative control. Three days post transfection, we amplified the target genomic site and found that the treatment with either PE-Cas9 or active Cas9, but not PE, led to a ~450-bp deletion amplicon. This amplicon was ~1-kb shorter than the amplicon without deletion (**Fig. 1c**). We digested the deletion amplicon with I-SceI endonuclease, and observed that only the PE-Cas9-treated group showed cut bands of expected size (~251bp and ~199bp), indicating insertion of the I-SceI recognition sequence (**Fig. 1d**). Using real-time quantitative PCR, we found that PE-Cas9 generates an accurate deletion-insertion frequency of 2.67±0.839% in total genomic DNA, whereas Cas9 seldom generated accurate deletion-insertion (0.0112±0.00717%, **Fig. 1e**). To further verify editing accuracy, we purified the deletion amplicon (~450-bp band in **Fig.1c**) and performed deep sequencing analysis. We found that PE-Cas9 mediates 27.0±1.83% accurate editing of total deletion events (**Fig. 1f**). Taken together, our findings suggest that, PEDAR outperforms prime editing and Cas9 editing in programming accurate large fragment deletion and simultaneous insertion.

PEDAR also generated unintended edits, classified as: (i) other deletion/insertion, including direct deletion without insertion and imperfect deletion-insertion, and (ii) small indels generated by individual pegRNA at the two cut sites, hereafter referred to as cut site_F and cut site_R. We measured the incidence of these events in total genomic DNA by real-time quantitative PCR, and observed that PE-Cas9 and Cas9 generated comparable rates of unintended edits (**Fig. 1e**). Next, we analyzed the deletion amplicons by deep sequencing. Of all the deletion events, PE-Cas9 generated 38.0±4.15% imperfect deletion-insertions caused by imprecise DNA repair or pegRNA scaffold insertion^17^ and a significantly lower rate of direct deletion without insertion than that mediated by active Cas9 (35.0±4.80% and 88.8±1.58%, respectively) (**Fig. 1f**). PE-Cas9-mediated unintended deletion edits with the highest sequencing reads are listed in **Supplementary Fig. 2a** and **Supplementary Table 3**. PE-Cas9 or Cas9 also introduced indels at the two cut sites without generating the desired deletion. Sanger sequencing of the amplicon without deletion (~1.4-kb band in **Fig.1c**) reveals no significant difference in small indels caused by PE-Cas9 and Cas9 (**Supplementary Fig. 2b**).

To explore the potential repair mechanism underlying PEDAR-mediated editing, we delivered PE-Cas9 with one pegRNA and one sgRNA targeting the *HEK3* locus into cells. And PE-Cas9 with paired pegRNAs serves as a positive control (**Supplementary Fig. 2c**). Although PE-Cas9 generated a ~450-bp deletion amplicon using one pegRNA and one sgRNA (**Supplementary Fig. 2d**), this amplicon failed to be digested into two distinct bands by I-Sce1 endonuclease (**Supplementary Fig. 2e**). Deep sequencing revealed that minimal accurate deletion-insertion (0.716±0.0868%) in the cells transfected with one pegRNA and one sgRNA, as compared to a 26.5±1.12% accurate editing rate in the cells treated with PEDAR (**Supplementary Fig. 2f**). This result demonstrates that the reverse-complementary sequences introduced by paired pegRNAs at the two cut sites are essential for directing accurate repair, resembling the annealing and ligation process in the MMEJ or SSA repair pathway^25,26^.

We also investigated how design of the pegRNAs, namely the length of the primer binding site (PBS) and the design of RT template, might affect editing efficiency of PEDAR. Our original PEDAR system used paired pegRNAs with 13-nt PBS. We designed two additional paired pegRNAs with 10-nt or 25-nt PBS targeting the *HEK3* locus as comparisons. Although all paired pegRNAs supported ~1-kb deletion (**Supplementary Fig. 3a**) and simultaneous insertion of the I-Sce1 recognition sequence (**Supplementary Fig. 3b**), the shorter and longer PBS lengths significantly impaired the accurate editing rate identified by deep sequencing (**Supplementary Fig. 3c**). To determine the effect of RT template design on editing efficiency, we designed an alternative pegRNA (pegRNA_alt) – similar to the pegRNA used in PE2^17^–by extending the RT template with a 14-nt sequence homologous to the region after the other cut site (**Supplementary Fig. 3d**). After transfecting the newly-designed paired pegRNAs with PE or PE-Cas9 into cells, we identified a deletion amplicon of the expected size (**Supplementary Fig. 3e**), and insertion of I-Sce1 recognition sequence was detected in the deletion amplicon (**Supplementary Fig. 3f**). Deep sequencing of the deletion amplicon reveals that pegRNA_alt significantly decreased PE-Cas9-mediated accurate editing rate compared to the original pegRNAs (**Supplementary Fig. 3g**). Surprisingly, co-transfection of PE and pegRNA_alt greatly improved the purity of deletion product (85.9±0.644% accurate editing in deletion amplicon, **Supplementary Fig. 3g**). However, the absolute accurate editing rate in total genomic DNA was comparable between PE/pegRNA_alt and PE-Cas9/pegRNA groups (**Supplementary Fig. 3h**), potentially due to the limited ability of Cas9 nickase to introduce larger deletion^23, 24^. Based on the collective findings, we elected to use pegRNAs with a 13-nt PBS and an RT template without adding the sequence homologous to target site after incision in the subsequent studies.

To assess the efficiency of PEDAR-mediated deletion-insertion at endogenous locus other than *HEK3* site, we targeted *DYRK1* locus for deleting a 995-bp DNA fragment and simultaneously inserting I-Sce1 recognition sequence. Treatment of HEK293T cells with PEDAR could lead to a ~507-bp deletion band (**Supplementary Fig. 4a**), and this amplified product could be digested by I-Sce1 endonuclease (**Supplementary Fig. 4b**). Deep sequencing of the deletion amplicon identified a 2.18±0.552% accurate editing efficiency (**Supplementary Fig. 4c**). We reasoned that the low G/C contents at the primer binding sequences of the two pegRNAs targeting *DYRK1* locus (23% of pegF and 31% of pegR) restricted the integration of the desired DNA fragment, which is consistent with a report showing poor PE efficiency when the GC content in PBS is less than 30%^27^.

To further understand the robustness of the PEDAR system, we explored its limits with respect to insertion size and deletion size. First, we set out to insert the I-Sce1 recognition sequence together with either Flag epitope tag (44bp total) or Cre recombinase *LoxP* site (60bp total) into the *HEK3* locus after deletion of a ~1-kb DNA fragment. Two paired pegRNAs were designed with either a 44-nt RT template or a 60-nt RT template, and the pegRNAs with 18-nt RT template serve for comparison (**Fig. 2a**). For all paired pegRNAs, PE-Cas9 generated the expected deletion (**Fig. 2b**) and inserted the desired sequence (**Fig.2c**) at the target site in cells. Deep sequencing revealed 13.7±1.51% (44bp-insertion) and 12.4±2.88% (60bp-insertion) accurate deletion-insertion rates within total deletion edits, which are significantly lower than the 22.6±0.267% accurate editing efficiency of PE-Cas9 when inserting a shorter sequence (18bp) (**Fig.2d**). To investigate the maximum deletion size generated by PEDAR, we designed two distinct paired pegRNAs with an offset of ~8kb or ~10kb to target the *CDC42* locus (**Fig.2e**). Using the indicated primers to amplify the corresponding target site, we observed the expected deletion amplicon (**Fig.2f**). After I-Sce1 endonuclease treatment, two digested bands were detected in the PE-Cas9-treated group (**Fig.2g**). Deep sequencing revealed 18.4±2.07% (8kb-del/18bp-ins) and 6.97±1.00% (10kb-del/18bp-ins) accurate deletion-insertion rates within the deletion amplicon (**Fig.2h**). In all, these data demonstrate the robustness and flexibility of PE-Cas9 in generating >10-kb larger deletion and up to 60-bp insertion.

**Fig 2.**
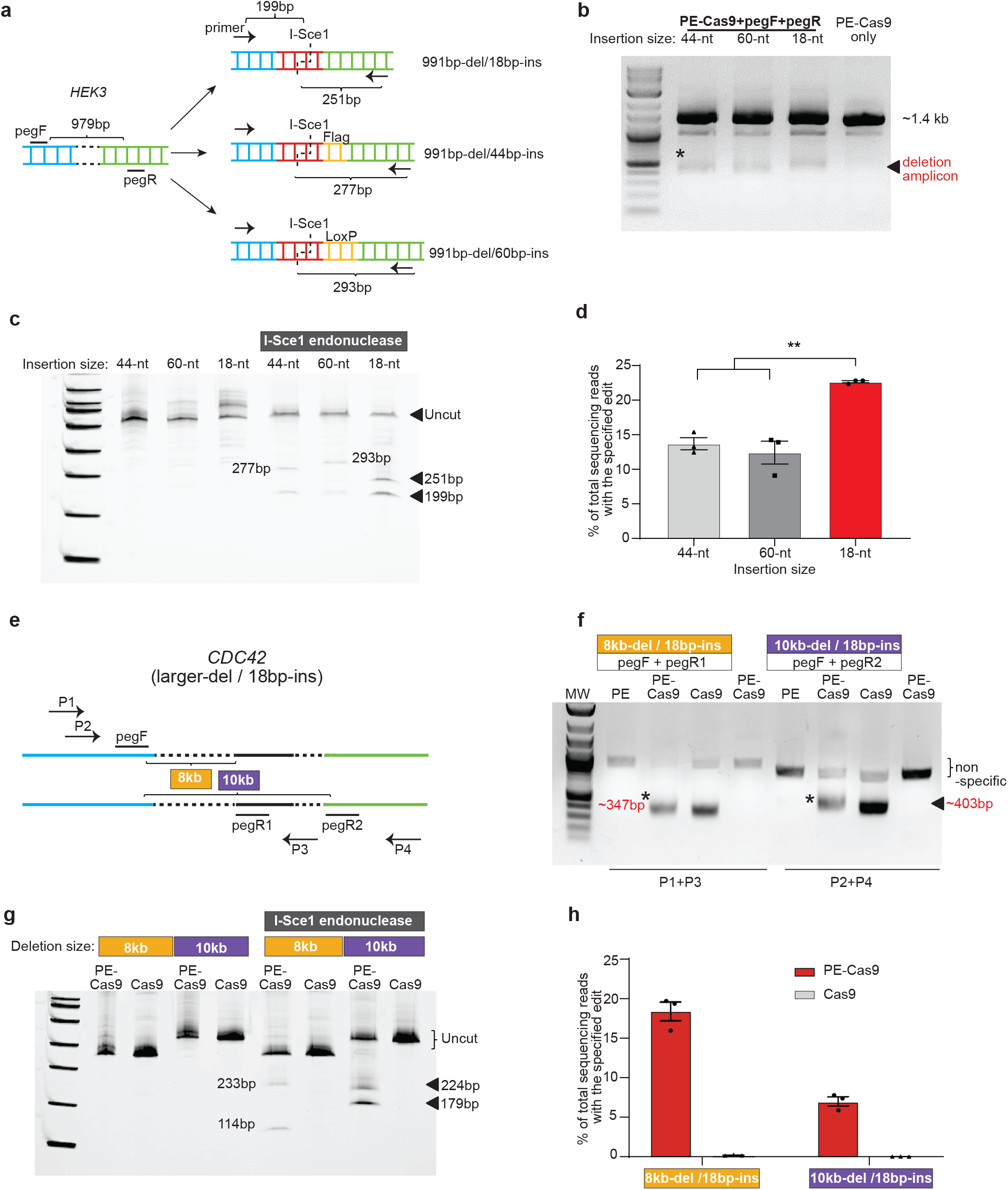
Flexibility of PEDAR in programming larger deletion and insertion in HEK293T cells. **a,** Insert DNA fragments of variable lengths (18-bp, 44-bp, and 60-bp) to the target site at *HEK3* locus. pegRNAs and primers for amplifying the target site are as shown. The expected sizes of digestion products after I-Sce1 treatment are shown. **b,** Amplification of target genomic region using primers spanning the cut sites at *HEK3* locus. HEK293T cells were transfected with PE-Cas9 and paired pegRNAs. The deletion amplicons are denoted with *. Cells transfected with PE-Cas9 alone serves as negative control. **c,** Deletion amplicons from groups shown in (b) were incubated with or without I-SceI endonuclease and analyzed in 4-20% TBE gel. Digested products are marked by arrows with expected sizes. The original amplicon is marked as “uncut”. **d,** Deep sequencing of deletion amplicons shown in (b). Bar chart shows accurate deletion-insertion rate. Data represent mean± SEM (n = 3 biologically independent samples). P=0.0024 (**), two-tailed t-test. **e,** Test the efficiency of PEDAR in mediating larger deletions. Paired pegRNAs spaced ~8-kb (pegF+pegR1) or 10-kb (pegF+pegR2) apart were designed as indicated to target the *CDC42* locus. Primers used to amplify the target genomic regions are as marked (P1+P3 and P2+P4). **f,** Target genomic region was amplified using the primers indicated in (e). Dual pegRNAs were transfected into HEK293T cells with PE, Cas9, or PE-Cas9. Cells transfected with PE-Cas9 alone serve as negative control. The deletion amplicons are marked with expected sizes (denoted with *). **g,** Deletion amplicons from Cas9- or PE-Cas9-treated groups shown in (f) were incubated with or without I-SceI endonuclease and analyzed in 4-20% TBE gel. Digested products are marked with expected sizes. The original amplicon is marked as “uncut”. **h,** Deep sequencing of deletion amplicons shown in (f). Bar chart shows rate of accurate deletion-insertion events. Data represent mean± SEM (n = 3 biologically independent samples).

Next, we asked whether PEDAR could generate large in-frame deletions and accurately repair genomic coding regions to restore gene expression. To answer this question, we used a HEK293T traffic light reporter (TLR) cell line^28, 29^, which contains a GFP sequence with an insertion and an mCherry sequence separated by a T2A (2A self-cleaving peptides) sequence. The disrupted GFP sequence causes a frameshift that prevents mCherry expression (**Fig. 3a**). We hypothesized that PEDAR could restore mCherry signal by accurately deleting the disrupted GFP and T2A sequence (~800 bp in length). We designed two pegRNAs targeting the promoter region before the start codon of GFP and the site immediately after T2A, respectively. In this approach, part of the Kozak sequence and start codon are unintentionally deleted due to the restriction of the PAM sequence. However, we designed the RT template at the 3’ end of pegRNAs to encode the Kozak sequence and start codon to ensure their insertion into the target site by reverse transcription (**Fig. 3a**).

**Fig 3.**
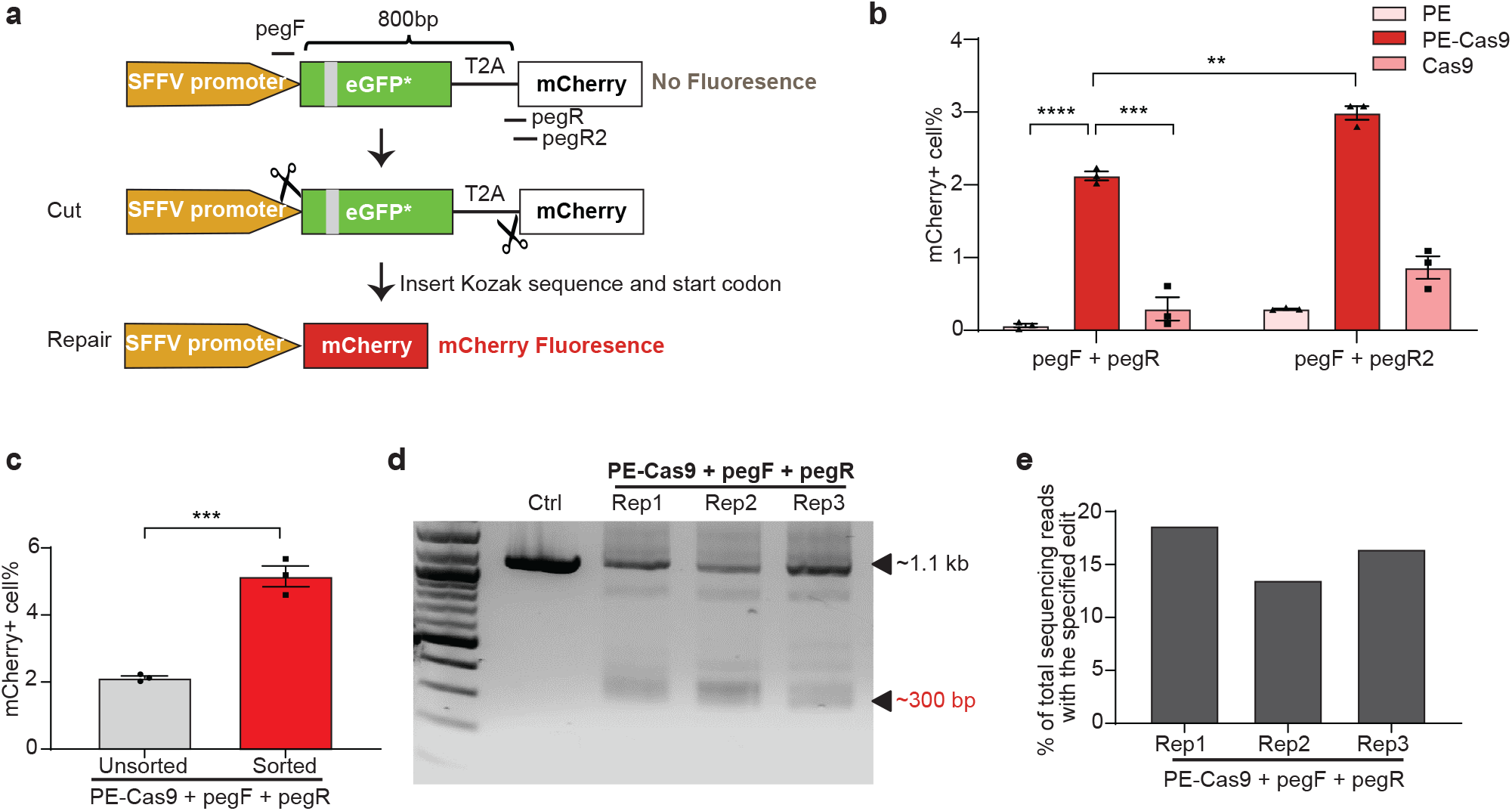
PEDAR generates in-frame deletion to restore mCherry expression in TLR reporter cells. **a,** Diagram of the TLR reporter system. GFP sequence is disrupted by an insertion (grey). Deleting the disrupted GFP sequence and inserting Kozak sequence and start codon will restore mCherry protein expression. **b,** TLR reporter cells transfected with indicated paired pegRNAs along with PE, PE-Cas9, or Cas9 were analyzed by flow cytometry, and the percentage of mCherry positive cells are shown among different groups. Data represent mean ± SEM (n = 3 biologically independent samples). P<0.0001 (****), =0.0004 (***), =0.0015 (**), two-tailed t-test. **c,** mCherry positive cell rate before and after sorting of cells with high transfection level. A plasmid expressing GFP was co-transfected with paired pegRNAs and PE-Cas9 into TLR cells. Three days later, cells with high GFP expression were selected for analyzing mCherry signal by flow cytometry. Data represent mean ± SEM (n = 3 biologically independent samples). P=0.0007 (***), two-tailed t-test. **d,** TLR reporter cells edited by PEDAR were selected by flow cytometry (for mCherry signal) and subjected to PCR amplification using primers spanning the two cut sites. The amplicon with the desired deletion is ~300 bp compared to a ~1.1-kb PCR products in control group. Rep: replicate; Ctrl: untreated TLR reporter cells. **e,** Efficiency of accurate deletion-insertion in three PEDAR-edited replicates (Rep 1-3) measured by deep sequencing of the deletion amplicons shown in (d).

We treated TLR reporter cells with dual pegRNAs (pegF+pegR) and either PE-Cas9, PE, or Cas9, and used flow cytometry to assess the mCherry signal. The frequency of mCherry positive cells was significantly higher in the PE-Cas9-treated group (2.12±0.105%) compared to PE- or Cas9-treated groups (**Fig. 3b** and **Supplementary Fig. 5a, b**). The mCherry positive cell rate was limited in all three replicates, likely because the cleavage efficiency of pegRNA at cut site_R (pegR) are very low (~1.8%; **Supplementary Fig. 5c**). Thus, we designed another pegRNA (pegR2) with a ~10.3% cleavage rate (**Fig. 3a** and **Supplementary Fig. 5c**) and assessed its efficiency in restoring mCherry expression. Indeed, the newly-designed paired pegRNAs significantly improved the mCherry positive cell rate (2.99±0.166%, **Fig. 3b** and **Supplementary Fig. 5d**). Alternatively, to enhance the editing rate, we explored the possibility of improving the expression level of gene editing agents in cells^30^. Co-transfection of cells with a fluorescent protein-expressing plasmid, followed by FACS sorting, would enrich for cells with high levels of transgene expression^31,32^. Thus, a GFP-expressing plasmid was co-transfected with PE-Cas9 and paired pegRNAs into TLR cells as an indicator of transfection efficiency. We observed a ~1.42-fold increase in mCherry positive cell rate after selection of cells with high GFP expression (**Fig. 3c** and **Supplementary Fig. 5e**). These results indicate that the editing efficiency of PEDAR largely relies on the efficiency of pegRNA and the expression level of gene editing components. To verify that PEDAR restored mCherry expression via accurate deletion-insertion, we sorted mCherry positive cells in PE-Cas9-treated groups and amplified the target sequence. In all three replicates, we detected a deletion amplicon that is ~800-bp shorter than the amplicon in untreated control cells (**Fig. 3d**). Further, we assessed the accurate editing rate by deep sequencing analysis of the ~300-bp deletion amplicon. The results revealed a 16.2±2.58% accurate deletion-insertion rate (**Fig. 3e**). The most common imperfect editing event across the three replicates restores mCherry open reading frame but the inserted sequence lacks three nucleotides compared to the intended insertion (**Supplementary Fig. 5f)**. These data demonstrate that PEDAR can repair genomic coding regions that are disrupted by large insertions.

Furthermore, to test the in vivo application of PEDAR, we utilized a Tyrosinemia I mouse model, referred to as Fah^ΔExon5^. This Tyrosinemia I model is derived by replacing a 19-bp sequence with a ~1.3-kb neo expression cassette^33^ at exon 5 of the *Fah* gene^34^ (**Fig. 4a**). This insertion disrupts the *Fah* gene to cause FAH protein deficiency and liver damage. To maintain body weight and survival, these mice are given water supplemented with NTBC [2-(2-nitro-4-trifluoromethylbenzoyl)-1,3-cyclohexanedione], a tyrosine catabolic pathway inhibitor. We hypothesized that PEDAR can correct the causative Fah^ΔExon5^ mutation by deleting the large insertion and simultaneously inserting the 19-bp fragment back to repair exon 5 (**Fig. 4b**). We engineered two pegRNAs targeting the genomic region before and after the inserted neo expression cassette, respectively. At the 3’ end of pegRNAs, a 22-bp RT template encoding the deletion fragment (19bp) plus a 3-bp sequence that is unintentionally deleted by PE-Cas9 was designed. PE-Cas9 and the two pegRNAs were delivered to the livers of mice (n=4) via hydrodynamic injection. Mice (n=2) treated with Cas9/pegRNAs serve as negative control. Mice were kept on NTBC water after treatment. One week later, the mice were euthanized, and immunochemical staining was performed on liver sections with FAH antibody. We detected FAH-expressing hepatocytes on PE-Cas9-treated liver sections (**Fig. 4c**), with a 0.76±0.25% correction rate (**Fig. 4d**). FAH expression was not detected in Cas9-treated mouse liver (**Fig. 4c**).

**Fig 4.**
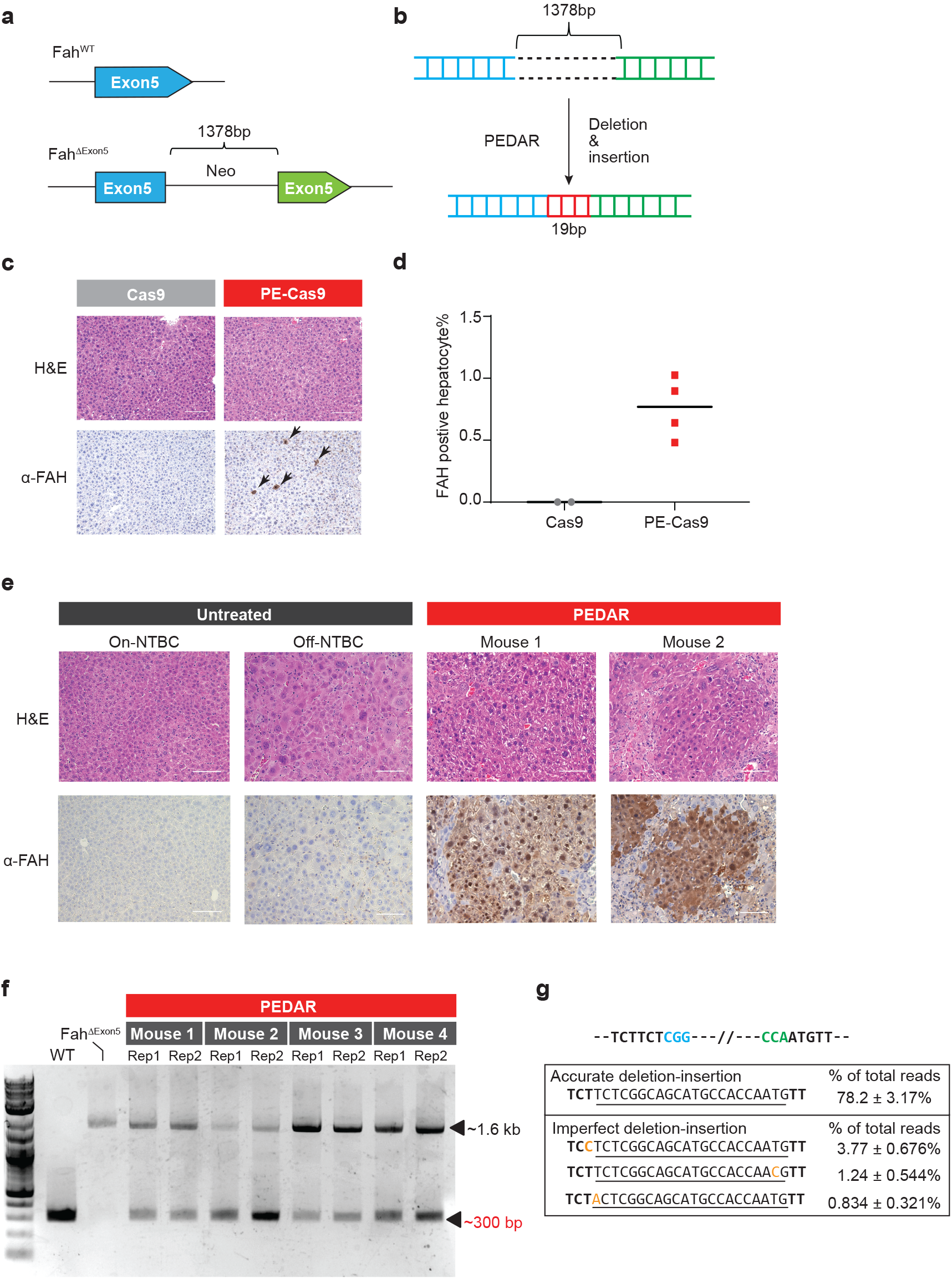
PEDAR corrects the pathogenic insertion in a Tyrosinemia I mouse model. **a,** The Tyrosinemia I mouse model, referred to as Fah^ΔExon5^, was derived by integrating a ~1.38-kb neo expression cassette at exon 5 of the *Fah* gene. **b,** Diagram showing the application of PEDAR to delete the ~1.38-kb insertion and concurrently repair the target region by inserting a 19-bp DNA fragment (marked in red). **c,** Immunohistochemistry staining and Hematoxylin and Eosin staining (H&E) of mouse liver sections seven days after injection of dual pegRNAs with Cas9 or PE-Cas9. FAH positive hepatocytes are pointed by arrows. Mice were kept on NTBC until being euthanized. Scale bar=100μm. **d,** Quantification of FAH expressing hepatocytes shown in (c). n=2 (Cas9-treated group), n=4 (PE-Cas9-treated group). **e,** Immunohistochemistry and H&E staining of mouse liver sections 40 days after injection of PE-Cas9 with dual pegRNAs. Mice (n=4) were kept off NTBC. Mouse 1 and 2 denote two representative mice from the treatment group. The liver sections from untreated Fah ^ΔExon5^ kept on or off NTBC serve as negative controls. Scale bar=100μm. **f,** Amplification of exon 5 of *Fah* gene from mouse livers 40 days post injection of PE-Cas9 and paired pegRNAs. The corrected amplicon size is around ~300 bp, compared to a ~1.6-kb amplicon without deletion. Four mice in treated group and two liver lobes (denoted as Rep 1 and 2) per mouse were analyzed. WT: wild type C57BL/6J mouse. Fah ^ΔExon5^: untreated Fah ^ΔExon5^ mouse. **g,** Accurate correction rate and the top-three imperfect editing events identified by deep sequencing. Two PAM sequences are in blue and green. The 22-bp intended insertion (19-bp deletion fragment plus a 3-bp unintentionally deleted sequence) is underlined. Mutated nucleotides in imperfect editing sequences are highlighted in yellow. Data represent mean ± SD (n = 8, two liver lobes/mouse; four mice in total).

Hepatocytes with corrected FAH protein will gain a growth advantage and eventually repopulate the liver^35^. Therefore, we delivered PE-Cas9 and the two pegRNAs via hydrodynamic injection to mice (n=4) and subsequently removed the NTBC supplement to allow repopulation. Untreated Fah^ΔExon5^ mice (on or off NTBC water) were used as controls. Forty days later, widespread FAH patches were observed in PE-Cas9-treated mouse liver sections, and the corrected hepatocytes showed normal morphology (**Fig. 4e, Supplementary Fig. 6a**). To understand the editing events in mouse liver, we amplified the target site by using PCR primers spanning exon 5, and identified the ~300-bp deletion amplicon in treated mice, indicating deletion of the ~1.3-kb insertion fragment (**Fig. 4f**). Deep sequencing of the ~300-bp deletion amplicon uncovered that accurate deletion-insertion constitutes 78.2±3.17% of total deletion events (**Fig. 4g**). We reasoned that, in this mouse model, hepatocytes with corrected FAH protein will outgrow cells with unintended editing, imposing a positive selection for desired editing. The average indel rates caused by each pegRNA at the *Fah* locus were 9.6% (cut site_F) and 0.14% (cut site_R) (**Supplementary Fig. 6b**). Although one mouse had a much higher average indel rate (27.7%) at cut site_F (Mouse 1 in **Supplementary Fig. 6b**), it did not negatively affect FAH protein expression (Mouse 1 in **Fig. 4e**). Overall, our data demonstrate the potential of using PEDAR in vivo to repair pathogenic mutations caused by large insertions.

Here, we expanded the application scope of prime editing by developing a PE-Cas9-based deletion and repair method – PEDAR – that can correct mutations caused by larger genomic rearrangements. Based on the design of prime editor, we combined Cas9 nuclease with an RT to engineer PE-Cas9. When delivered with paired pegRNAs, PE-Cas9 couples target deletion and insertion to direct the desired genome edits. Our PEDAR system is similar to a recently developed paired prime editing method, called *PRIME-Del*, that can introduce 20- to 700-bp target deletions and up to 30-bp insertions^36^. Compared to *PRIME-Del*, PEDAR seems to be more error-prone, introducing higher fractions of direct deletion and imperfect deletion-insertion (**Supplementary Fig. 3g**); however, both editors exhibited comparable absolute accurate rates in total genomic DNA (**Supplementary Fig. 3h**). Importantly, we show that PEDAR is able to introduce >10-kb target deletions and up to 60-bp insertions in cells, both of which are larger than what primer editors can generate^17, 36^. Moreover, PEDAR can program target deletion-insertion editing in quiescent hepatocytes in mouse liver, where HDR is not favorable^37^. Thus, PEDAR is a robust genome editing technique to couple large deletion and desired insertion in vitro and in vivo.

Despite the relative editing efficiency and accuracy of PE-Cas9 being higher than PE and Cas9, the overall efficiency of PEDAR was limited. In this study, we found that PEDAR editing was restricted by both cleavage activity of pegRNAs and transfection efficiency. Moreover, we revealed that PBS length and RT template length affect editing efficiency. Thus, we recommend that multiple pegRNA sequences with distinct spacer sequences, PBSs, or RT templates should be designed and compared to maximize PEDAR efficiency. Lastly, we show that PEDAR might share a similar mechanism with MMEJ or SSA during repairing the DSB. Thus, we expect that treatment with an MMEJ or SSA enhancer^38, 39^ could further improve the efficiency of PEDAR editing. Further studies are needed to identify the exact pathway or component that is involved in the repair process^25^. Although current studies claim that prime editing shows minimum off-target editing^17, 40, 41^, its genome-wide off-target effects in mammalian cells are not fully studied. Moreover, active Cas9 is known to bind unintended genomic sites for cleavage, introducing genome-wide off-target editing^42^. Thus, further evaluation of potential off-target effects by PEDAR at the whole genome level should be performed.

Finally, we propose that PEDAR could also be used to correct genome duplications (**Supplementary Fig. 7a**), which constitute ~10% of all human pathogenic mutations according to the ClinVar database^1^. One such genome duplication of high clinical significance is the trinucleotide CAG repeat expansion in the HTT gene – the root cause of Huntington disease^43^. Future studies should investigate whether PEDAR could accurately remove this expansion to reduce CAG repeat length. Thus, our findings have potential implications for the gene therapy field. The significance of PEDAR also extends to basic biology, where it could be used for protein function studies (**Supplementary Fig. 7b**). Previous studies introduce in-frame deletions by a “tiling CRISPR” method to explore the functional domain of specific genomic-coding or long non-coding regions^44, 45^. Here, we demonstrate that PEDAR exhibits a higher efficiency in mediating in-frame deletion compared to the canonical CRISPR/Cas9 system.

In summary, PEDAR offers a novel genetic tool for mediating precise large target genomic deletion and simultaneous insertion.

## Methods

### Cell Culture and Transfection

Human embryonic kidney (HEK293T) cells (ATCC) and HEK293T-TLR cells^28, 29^ were maintained in Dulbecco’s Modified Eagle’s Medium (Corning) supplemented with 10% fetal bovine serum (Gibco) and 1% Penicillin/ Streptomycin (Gibco). Cells were seeded at 70% confluence in 12-well cell culture plate one day before transfection. 1.5 μg PE-Cas9, and 1 μg paired pegRNAs (0.5 μg each) was transfected with Lipofectamine 3000 reagent (Invitrogen).

### PegRNA design and clone

Sequences for pegRNAs are listed in **Supplementary Table 1**. Plasmids expressing pegRNAs were constructed by Gibson assembly using BsaI-digested acceptor plasmid (Addgene #132777) as the vector.

### Mouse experiments

All animal study protocols were approved by the UMass Medical School IACUC. Fah ^ΔExon5^ mice^34^ were kept on 10mg/L NTBC water. 30μg PE-Cas9 or Cas9 plasmid and 15μg paired pegRNA expressing plasmids were injected into 9-week-old mice. One week later, NTBC supplemented water was replaced with normal water, and mouse weight was measured every two days. As per our guidelines, when the mouse lost 20% of its body weight relative to the first day of measurement (day when NTBC water was removed), the mouse was supplemented with NTBC water until the original body weight was achieved. After 40 days, mice were euthanized according to guidelines.

### Immunohistochemistry

Portion of livers were fixed with 4% formalin, embedded in paraffin, sectioned at 5 μm and stained with hematoxylin and eosin (H&E) for pathology. Liver sections were de-waxed, rehydrated, and stained using standard immunohistochemistry protocols^46^. The following antibody was used: anti-FAH (Abcam, 1:400). The images were captured using Leica DMi8 microscopy.

### Genomic DNA extraction, amplification, and digestion

To extract genomic DNA, HEK293T cells (3 days post transfection) were washed with PBS, pelleted, and lysed with 50μl Quick extraction buffer (Epicenter) and incubated in a thermocycler (65°C 15 min, and 98°C 5 min). PureLink Genomic DNA Mini Kit (Thermo Fisher) was used to extract genomic DNA from two different liver lobes (~10 mg each) per mouse. Target sequences were amplified using Phusion Flash PCR Master Mix (Thermo Fisher) with the primers listed in **Supplementary Table 2**. PCR products were analyzed by electrophoresis in a 1% agarose gel, and target amplicons were extracted using DNA extraction kit (Qiagen). 10 ng of purified PCR products were incubated with I-SceI endonuclease (NEB) according to manufacturer’s instruction. One-hour post incubation, the product was visualized and analyzed by electrophoresis in 4-20% TBE gel (Thermo).

### Tracking of Indels by Decomposition (TIDE) analysis to calculate indel rates at two cut sites

The sequences around the two cut sites of the target locus were amplified using Phusion Flash PCR Master Mix (Thermo Fisher) with the primers as listed in **Supplementary Table 2**. Sanger sequencing was performed to sequence the purified PCR products, and the trace sequences were analyzed using TIDE software (https://tide.nki.nl/). The alignment window of left boundary was set at 10-bp.

### Quantification of total genomic DNA to determine absolute editing rate of PEDAR

Real-time quantitative PCR (qPCR) was used to calculate the absolute editing rate in total genomic DNA at the *HEK3* locus. Quantitative PCR was performed with SsoFast EvaGreen Supermix (Bio-rad). Primers within the deletion region (P1 and P2), spanning the deletion region (P3 and P4), or across the deletion-insertion junction (P5 and P6) were designed (**Supplementary Fig. 8a**). Two 250-bp DNA fragments (referred to as WT and Edited) of the same sequence with unedited or accurately edited target site were designed and serially diluted, serving as standard templates (**Supplementary Fig. 8b**). Using indicated primers and templates to perform quantitative PCR, three standard curves were generated, reflecting the correlation between qPCR cycle number and the concentration of DNA without 991-bp deletion (**Supplementary Fig. 8c**), with 991-bp deletion (**Supplementary Fig. 8d**), or with accurate 991-bp deletion/18-bp insertion (**Supplementary Fig. 8e**). Finally, three rounds of quantitative PCR were performed using the edited genomic DNA as template and corresponding primer pairs (P1+P2, P3+P4, or P5+P6). The standard curves were applied to calculate the absolute copy number of genomic DNA with deletion, without deletion, or with accurate deletion-insertion.

The absolute rates of each type of editing introduced by PEDAR were calculated as follows: (1) Accurate deletion-insertion editing rate = copy number of DNA with accurate deletion-insertion / copy number of DNA with and without deletion. (2) Other deletion-insertion rate = (copy number of DNA with deletion - copy number of DNA with accurate deletion-insertion) / copy number of DNA with and without deletion. (3) Absolute rate of small indels at two cut sites = copy number of DNA without deletion x indel rate at distinct cut site calculated by TIDE / copy number of DNA with and without deletion

### Flow Cytometry analysis

To assess mCherry recovery rate, post-editing HEK293T-TLR cells were trypsinized and analyzed using the MACSQuant VYB Flow Cytometer. Untreated HEK293T-TLR cells were used as a negative control for gating. To select cells with high transfection efficiency, 0.25 μg GFP plasmid was co-transfected with PE-Cas9 and paired pegRNAs into TLR cells. Three days post transfection, cells were trypsinized and analyzed using the MACSQuant VYB Flow Cytometer. Cells transfected with GFP plasmid alone were used as a negative control for gating. Cells with high expression level of GFP (~20% of total population) were selected for analyzing mCherry signal. All data were analyzed by FlowJo10.0 software.

### High throughput DNA sequencing of genomic DNA samples

Genomic sites of interest were amplified from genomic DNA using specific primers containing illumina forward and reverse adaptors (listed in **Supplementary Table 2**). To quantify the percentage of desired deletion-insertion by PE-Cas9 or Cas9, we amplified the fragment containing deletions (~200 bp in length) from total genomic DNA to exclude length-dependent bias during PCR amplification. 20 μL PCR1 reactions were performed with 0.5 μM each of forward and reverse primer, 1 μL of genomic DNA extract or 300ng purified genomic DNA, and 10 μL of Phusion Flash PCR Master Mix (Thermo Fisher). PCR reactions were carried out as follows: 98°C for 10s, then 20 cycles of [98°C for 1 s, 55°C for 5 s, and 72°C for 10 s], followed by a final 72°C extension for 3 min. After the first round of PCR, unique Illumina barcoding reverse primer was added to each sample in a secondary PCR reaction (PCR 2). Specifically, 20 μL of a PCR reaction contained 0.5 μM of unique reverse Illumina barcoding primer pair and 0.5 μM common forward Illumina barcoding primer, 1 μL of unpurified PCR 1 reaction mixture, and 10 μL of Phusion Flash PCR Master Mix. The barcoding PCR2 reactions were carried out as follows: 98 °C for 10s, then 20 cycles of [98°C for 1 s, 60°C for 5 s, and 72°C for 10 s], followed by a final 72 °C extension for 3 min. PCR 2 products were purified by 1% agarose gel using a QIAquick Gel Extraction Kit (Qiagen), eluting with 15 μL of Elution Buffer. DNA concentration was measured by Bioanalyzer and sequenced on an Illumina MiSeq instrument (150bp, paired-end) according to the manufacturer’s protocols. Paired-end reads were merged with FLASh^47^ with maximum overlap length equal to 150 bp. Alignment of amplicon sequence to the reference sequence was performed using CRISPResso2^48^. To quantify accurate deletion-insertion edits, CRISPResso2 was run in HDR mode using the sequence with desired deletion-insertion editing as the reference sequence. The editing window is set to 15-bp. Editing yield was calculated as: [# of HDR aligned reads] ÷ [total aligned reads].

### ClinVar data analysis

The ClinVar variant summary was obtained from NCBI ClinVar database (accessed Dec 31,2020). Variants with pathogenic significance were filtered by allele ID to remove duplicates. All pathogenic variants were categorized according to mutation type. The fractions of distinct mutation types were calculated using GraphPad Prism8.

## Data Availability

A reporting summary for this article is available as a Supplementary Information file. All other data are available from the corresponding author upon reasonable request.

## Acknowledgements

We thank C. Mello, P. Zamore, S. Wolfe, T. Flotte, and E. Sontheimer for discussions and E. Haberlin for editing the manuscript. We thank Dr. Erik Sontheimer (UMass Medical School) for providing the HEK293T-TLR cell line and Dr. Markus Grompe (Oregon Health & Science University) for providing the Fah ^ΔExon5^ mice. We thank Y. Liu, Yuehua Gu, and E. Kittler in the UMass Morphology, Flow Cytometry, and Deep Sequencing Cores for support. W.X was supported by grants from the National Institutes of Health (DP2HL137167, P01HL131471 and UG3HL147367), American Cancer Society (129056-RSG-16-093), the Lung Cancer Research Foundation, and the Cystic Fibrosis Foundation. T.J was supported by grants from National Institutes of Health (K99HL153940).

## Competing Interests

UMass has filed a patent application on this work. W.X. is a consultant for the Cystic Fibrosis Foundation Therapeutics Lab. The other authors declare no competing interests.

## Author Contributions

T.J and W.X designed the study. T.J performed experiments. T.J, X-O.Z, and Z.W analyzed data. T.J and W.X wrote the manuscript with comments from all authors.

---

**Supplementary Fig. 1.**
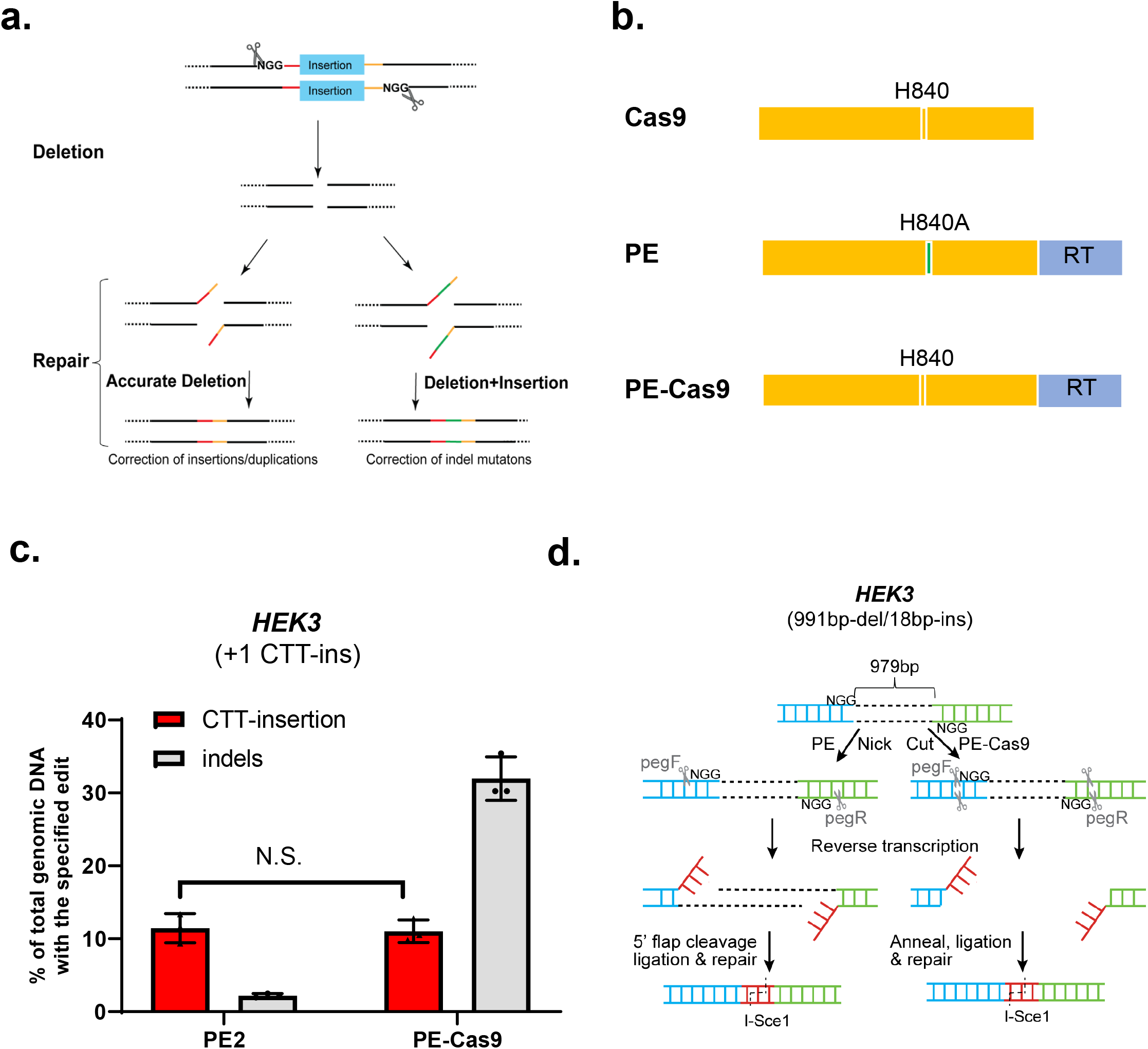
**a,** Proposed model of CRISPR-associated gene correction of pathogenic mutations caused by insertions/duplications or indels. The pathogenic insertion is removed by CRISPR under the guidance of dual sgRNAs targeting two complementary strands of DNA, while the repair or insertion is concurrently performed at the cut site. **b,** Engineering of PE-Cas9. PE-Cas9 is engineered by replacing the Cas9^H840A^ in PE with an active Cas9 nuclease. **c,** Comparison of PE2- and PE-Cas9-mediated insertion of a 3-bp nucleotide sequence (“CTT”) at the nicking/cut site of the *HEK3* locus. HEK293T cells were transfected with a pegRNA and PE-Cas9 or PE. The rate of accurate insertion and indels was assessed by deep sequencing. P=0.7929 (Not significant, N.S.), two-tailed t-test. **d,** Diagram of concurrent deletion of a 991-bp DNA fragment and insertion of 18-bp I-SceI recognition sequence (red) by PE or PE-Cas9 and paired pegRNAs. Two pegRNAs with an offset of 979 bp (distance between the two ‘NGG’ PAM sequences) were designed and transfected with PE or PE-Cas9 into cells.

**Supplementary Fig. 2.**
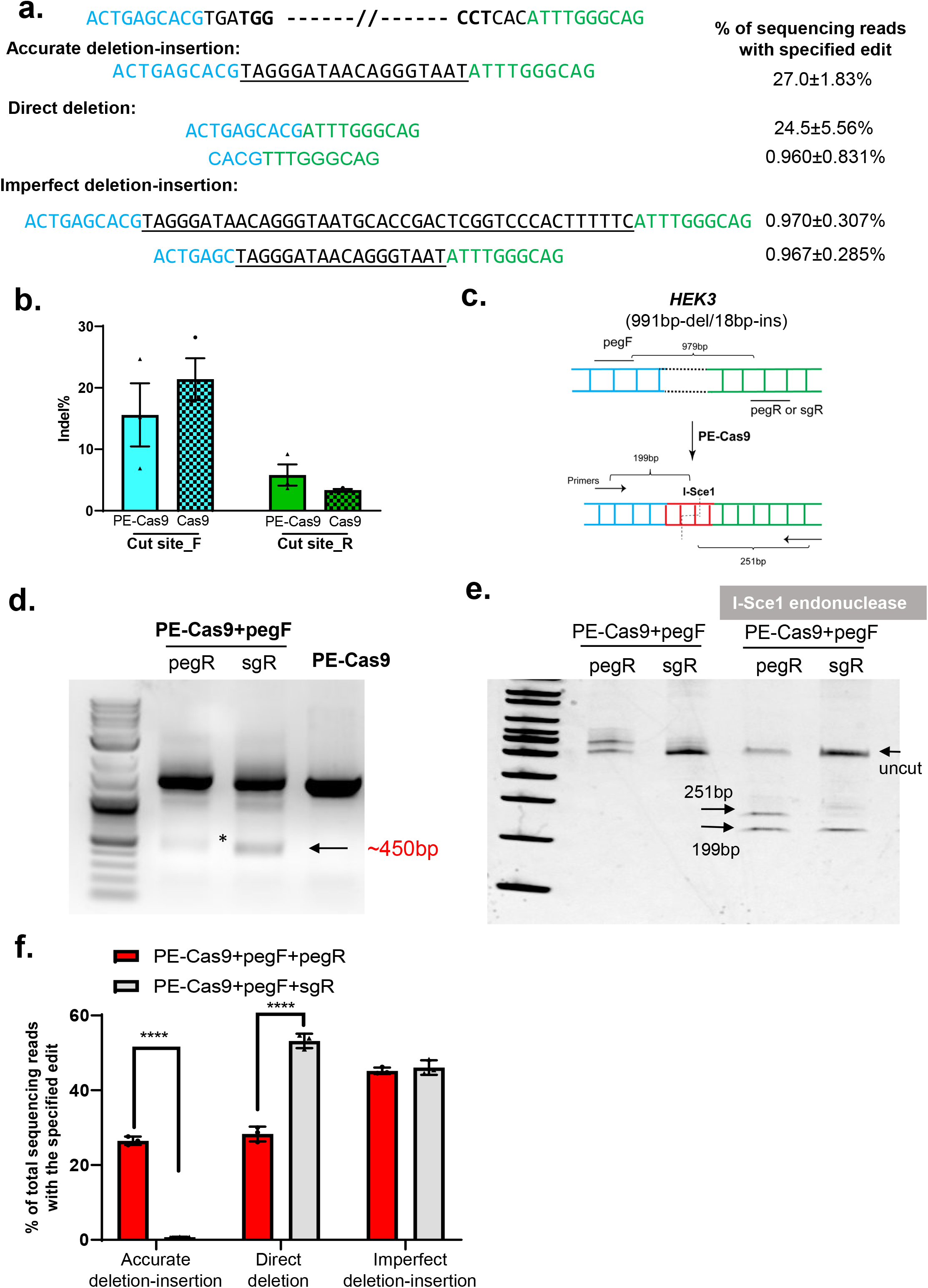
**a,** PE-Cas9-mediated editing events with highest reads across three replicates by deep sequencing. The two PAM sequences are in bold, and the original sequences before or after the two cut sites are highlighted in blue and green. The inserted sequence is underlined. Data represent mean ± SD (n =3 biologically independent samples). **b,** The indel rates generated by individual pegRNA at the two cut sites of *HEK3* site, hereafter referred to as cut site_F and cut site_R, assessed by sanger sequencing. Data represent mean ± SEM (n =3 biologically independent samples). **c,** Diagram showing that treatment of the edited PCR product with I-Sce1 endonuclease would lead to two DNA fragments of 199-bp and 251-bp at length. **d,** Amplification of target genomic region using primers that span the cut sites at *HEK3* locus. HEK293T cells were transfected with PE-Cas9, pegF, and pegR or sgR. The ~450-bp band is the deletion amplicon. Cells transfected with PE-Cas9 alone serve as negative control. **e,** Deletion amplicons from pegR or sgR-treated groups shown in (d) were incubated with or without I-SceI endonuclease and analyzed in 4-20% TBE gel. The digested products are marked by arrows with expected sizes. The original amplicon is marked as “uncut”. **f,** Deep sequencing of deletion amplicons shown in (d). Bar chart shows distribution of all the deletion events, including accurate deletion-insertion, direct deletion (deletion without any insertions), or imperfect deletion-insertion. Data represent mean ± SEM (n = 3 biologically independent samples). P=<0.0001 (****), two-tailed t-test.

**Supplementary Fig. 3.**
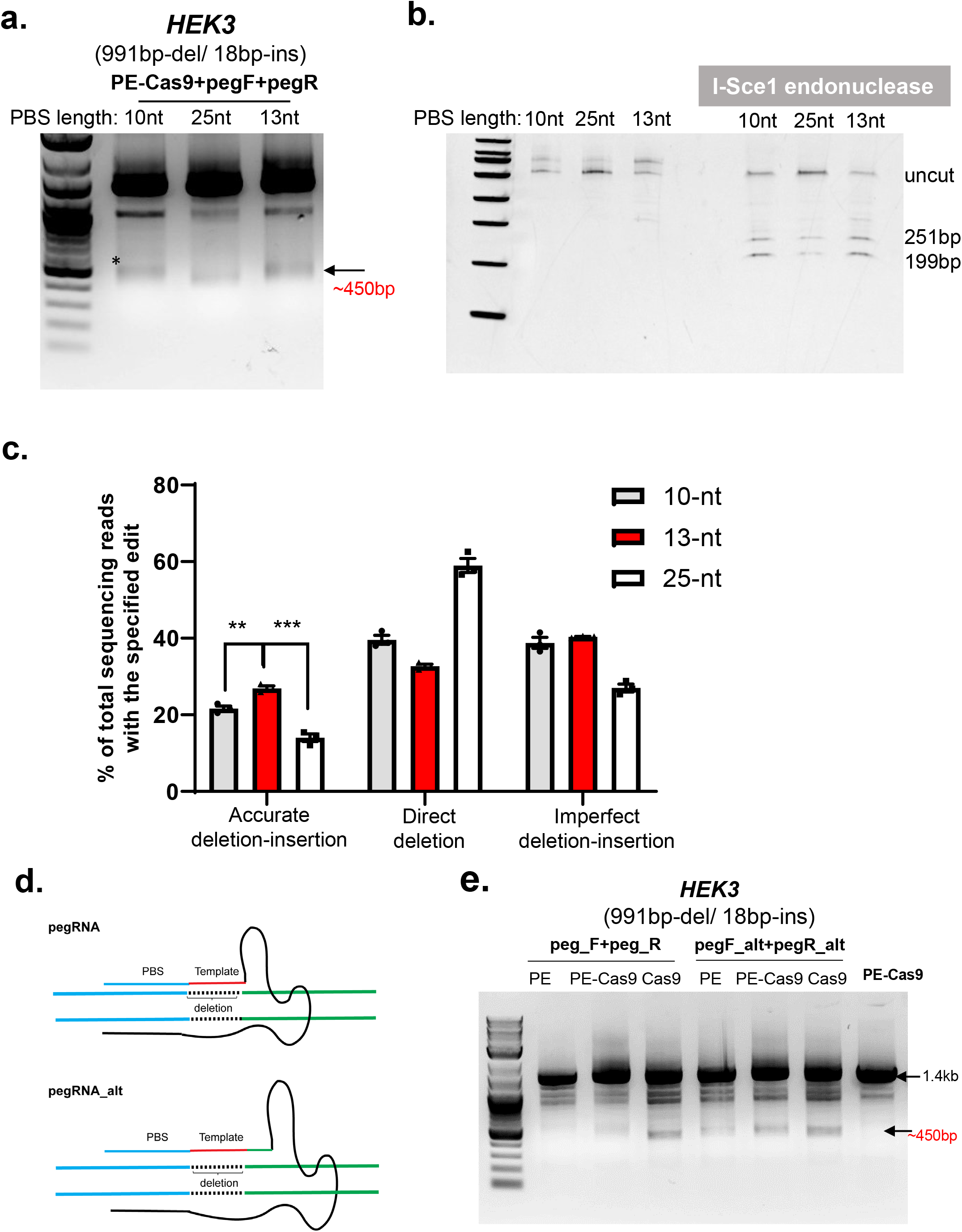
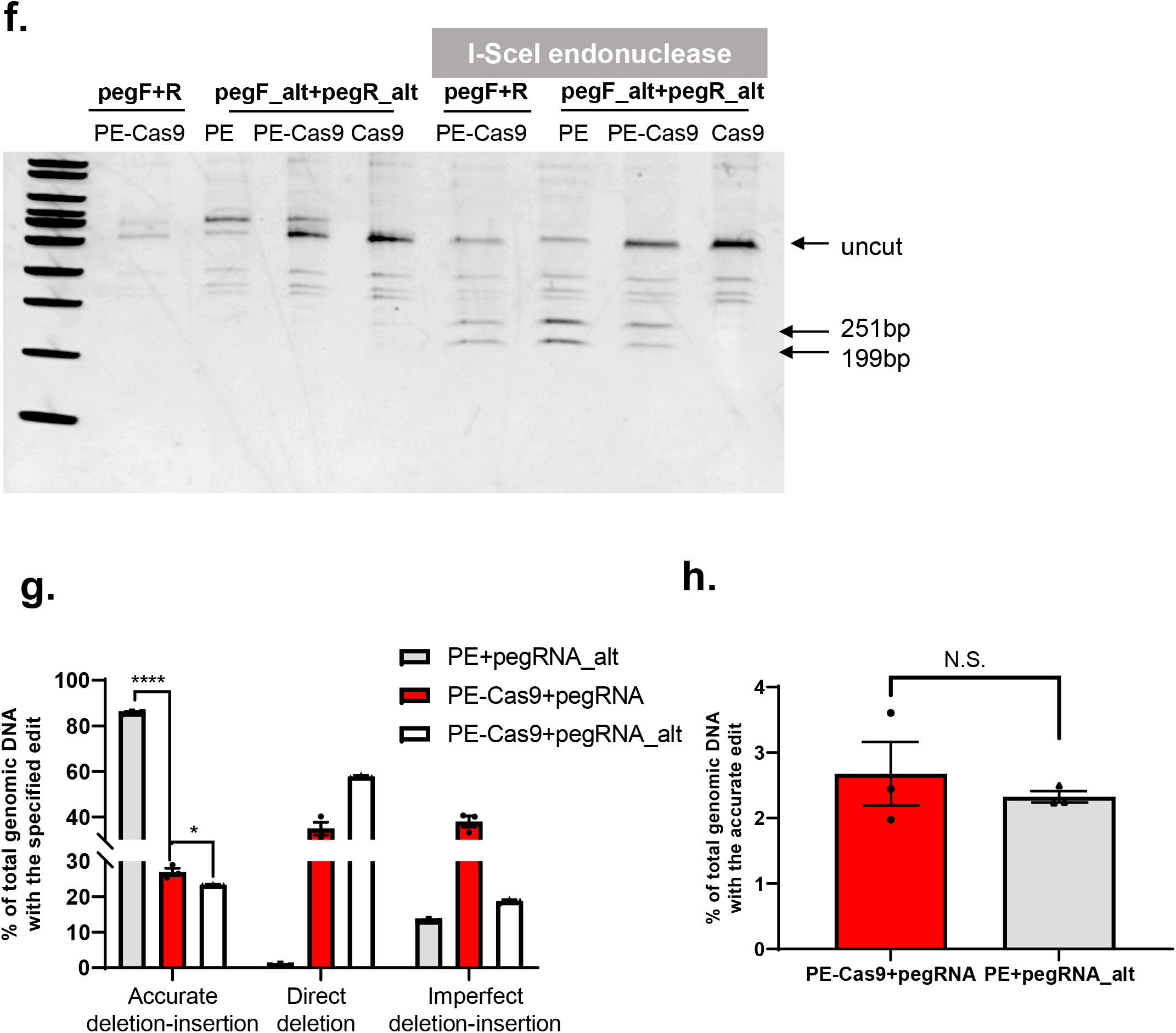
**a,** Amplification of target genomic region using primers that span the cut sites at *HEK3* site. Paired pegRNAs with indicated lengths of primer binding sequence were designed and transfected with PE-Cas9 into HEK293T cells. The ~450-bp band (denoted with *) is the expected deletion amplicon. **b,** Deletion amplicons from groups shown in (a) were incubated with or without I-SceI endonuclease and analyzed in 4-20% TBE gel. The digested products are marked with expected sizes. The original amplicon is marked as “uncut”. **c,** Deep sequencing of deletion amplicons shown in (a). Bar chart shows distribution of all the deletion events, including accurate deletion-insertion, direct deletion (deletion without any insertions), or imperfect deletion-insertion. Data represent mean± SEM (n = 3 biologically independent samples). P=0.0046 (**), =0.0004 (***), two-tailed t-test. **d,** Design alternative pegRNA (pegRNA_alt) by extending RT template with a 14-nt sequence homologous to the region after the other cut site. **e,** Amplification of target genomic region using primers that span the cut sites at the *HEK3* locus. HEK293T cells were transfected with Cas9, PE-Cas9 or PE along with paired pegRNAs as indicated. The ~450-bp band is the expected deletion amplicon. Cells transfected with PE-Cas9 alone serve as negative control. **f,** Deletion amplicons from groups shown in (e) were incubated with or without I-SceI endonuclease and analyzed in 4-20% TBE gel. The digested products are marked by arrows with expected sizes. The original amplicon is marked as “uncut”. **g,** Deep sequencing of deletion amplicons shown in (e). Bar chart shows distribution of all deletion events, including accurate deletion-insertion, direct deletion (deletion without any insertions), or imperfect deletion-insertion. Data represent mean ± SEM (n = 3 biologically independent samples). P=<0.0001 (****), =0.0269 (*), two-tailed t-test. **h,** Absolute accurate editing rates of PEDAR (PE-Cas9+pegRNA) and *PRIME-Del* (PE+pegRNA_alt) at the *HEK3* locus calculated by quantitative PCR in total genomic DNA. P=0.5156 (N.S., not significant).

**Supplementary Fig. 4.**
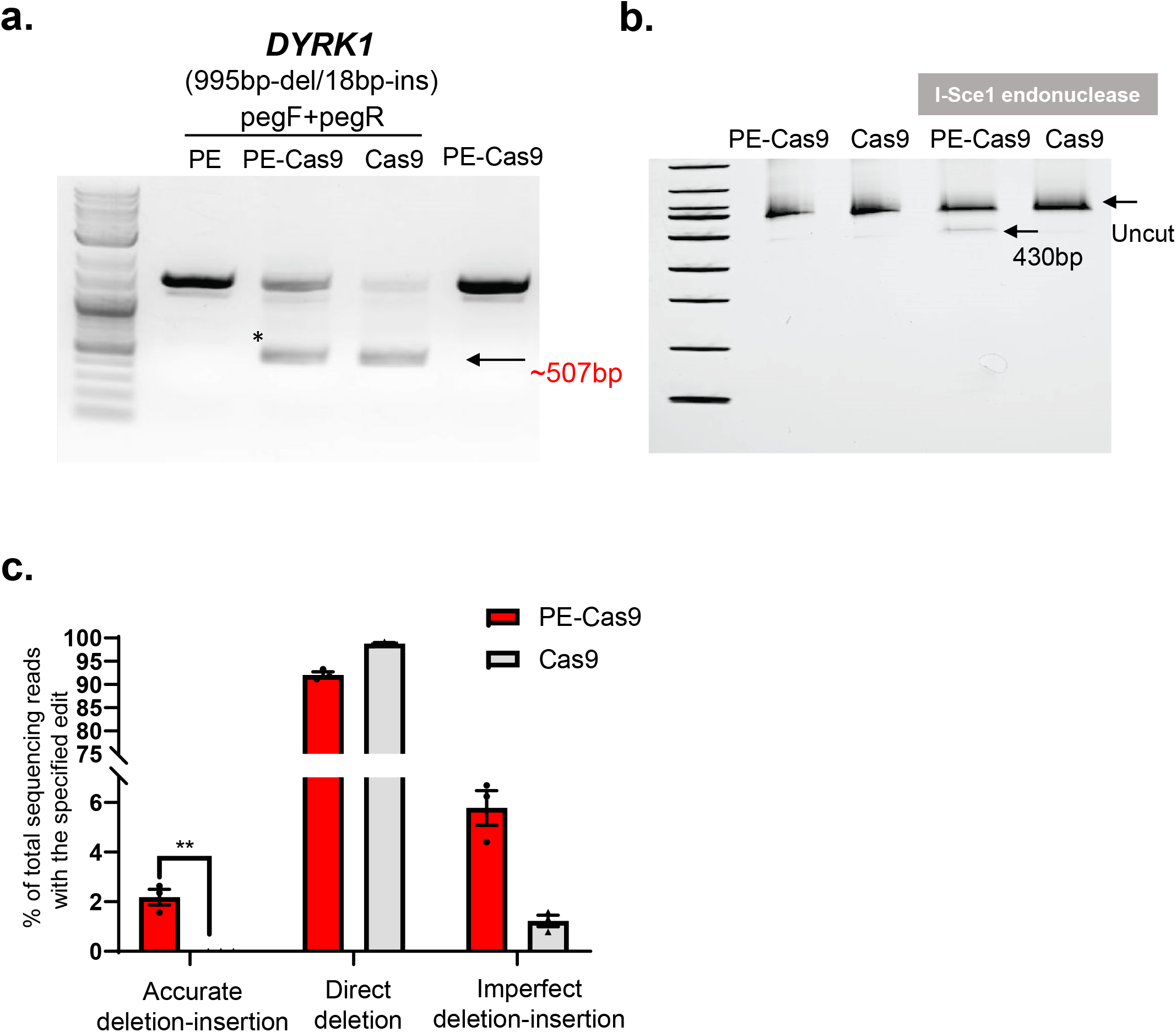
**a,** Amplification of target genomic region using primers that span the cut sites at *DYRK1* locus. The paired pegRNAs targeting complementary DNA strand are denoted as pegF and pegR. HEK293T cells were transfected with PE, Cas9, or PE-Cas9 with or without paired pegRNAs. The size of deletion amplicon (denoted with *) is indicated. **b,** Deletion amplicons from groups shown in (a) were incubated with or without I-SceI endonuclease and analyzed in 4-20% TBE gel. The digested products are marked by arrows with expected sizes. The original amplicon is marked as “uncut”. **c,** Deep sequencing of deletion amplicons shown in (a). Bar chart shows distribution of all the deletion events, including accurate deletion-insertion, direct deletion (deletion without any insertions), or imperfect deletion-insertion. Data represent mean ± SEM (n = 3 biologically independent samples). P=0.0036 (**), two-tailed t-test.

**Supplementary Fig. 5.**
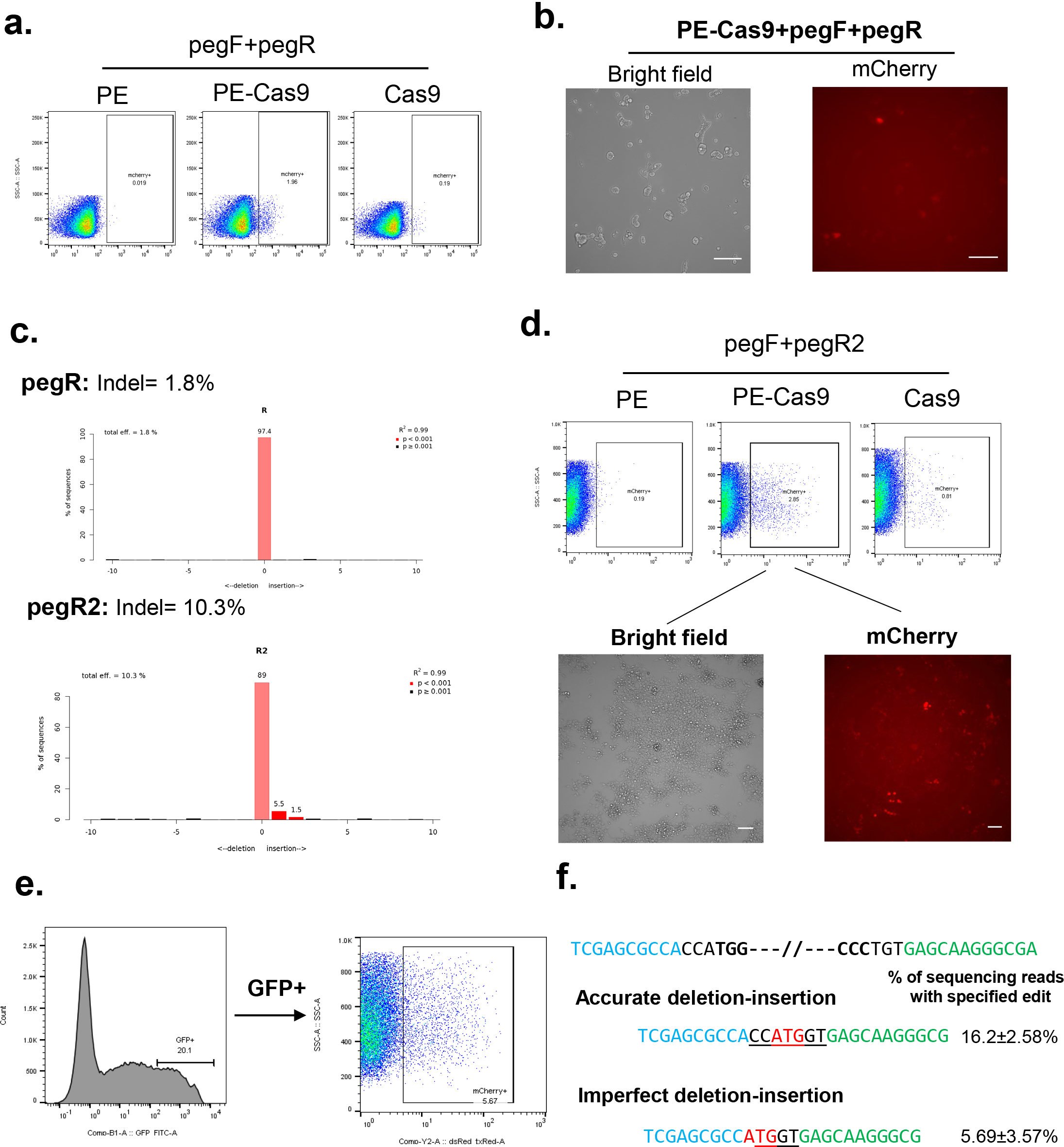
**a,** A representative flow cytometry plot shows the gating of mCherry positive cells in PE-, PE-Cas9-, or Cas9-treated groups. **b,** Image of mCherry positive cells treated by PEDAR. Scale bar =100 μm. **c,** TIDE results showing the indels introduced by two distinct pegRNAs (pegR and pegR2) at TLR locus. Cas9 was transfected together with pegR or pegR2 in HEK293T cells. Indel rates were analyzed by Tide software (http://tide.nki.nl). **d,** Upper panel: representative flow cytometry plot shows the gating of mCherry positive cells in PE, PE-Cas9, or Cas9-treated groups. Lower panel: image of mCherry positive cells treated by PEDAR. Scale bar =100 μm. **e,** Flow cytometry plots show the gating of TLR cells with high GFP expression (left panel; ~20% of total population) and the gating of mCherry positive cell after sorting out the GFP positive cells (right panel). GFP expression serves as an indicator of transfection rate. **f,** The rate of accurate editing and the most common imperfect deletion-insertion editing events identified across three replicates. The two PAM sequences are in bold, and the original sequences before or after the two cut sites are highlighted in blue and green. The inserted sequence is underlined. Start codon is highlighted in red. Data represent mean ± SD (n =3 biologically independent samples).

**Supplementary Fig. 6.**
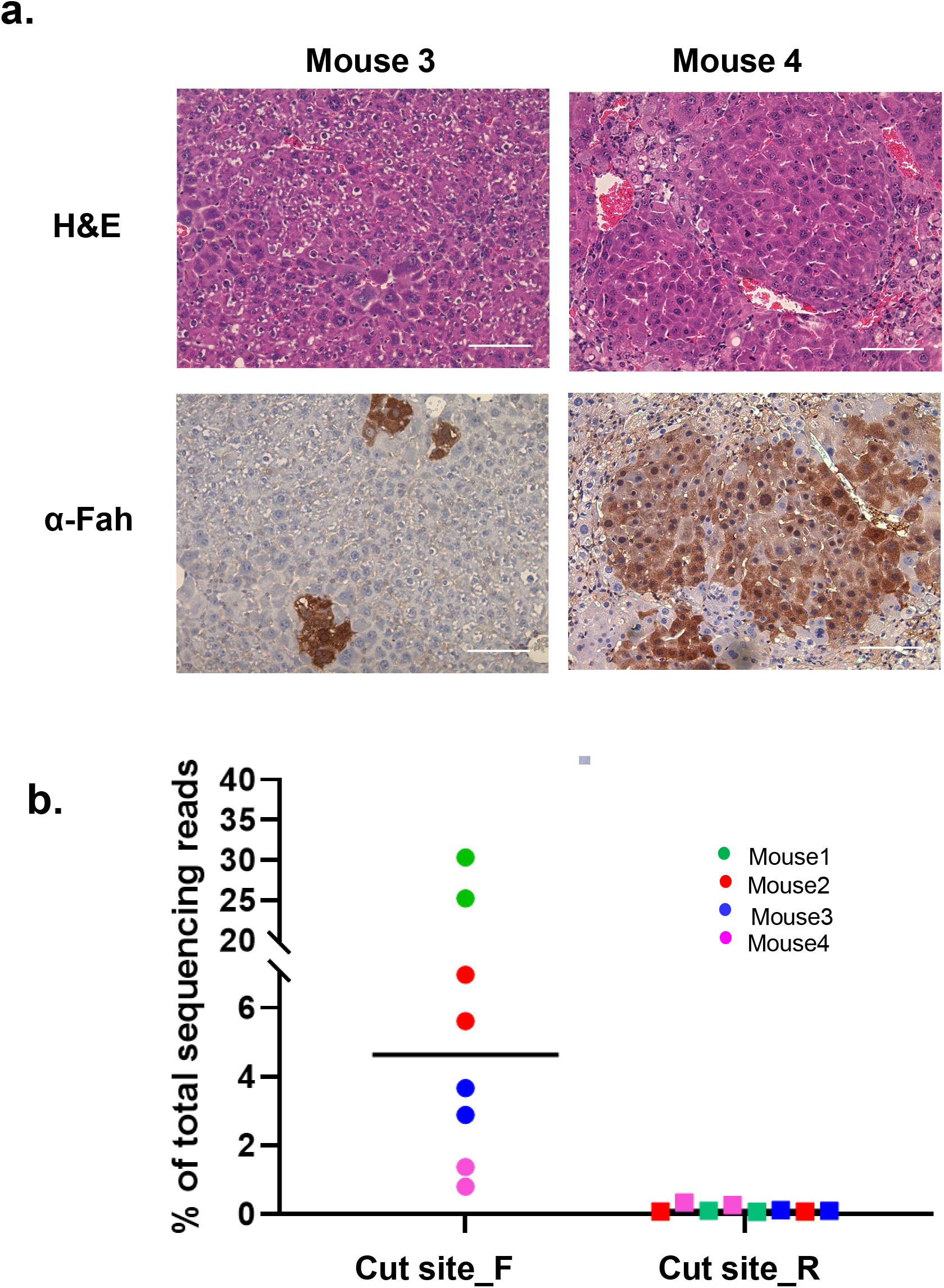
**a,** Immunohistochemistry staining and Hematoxylin and Eosin staining (H&E) of mouse liver sections 40 days after injection of PE-Cas9 with dual pegRNAs. Mice were kept off NTBC after treatment until euthanizing. Mouse 3 and 4 denote two distinct mice from the treatment group. Scale bar=100μm. **b,** Indel rates generated by individual pegRNA at the two cut sites at the *Fah* locus. Four mice in treated group and two liver lobes per mouse were analyzed. The dots with the same color indicate samples from two liver lobes of the same mouse.

**Supplementary Fig. 7.**
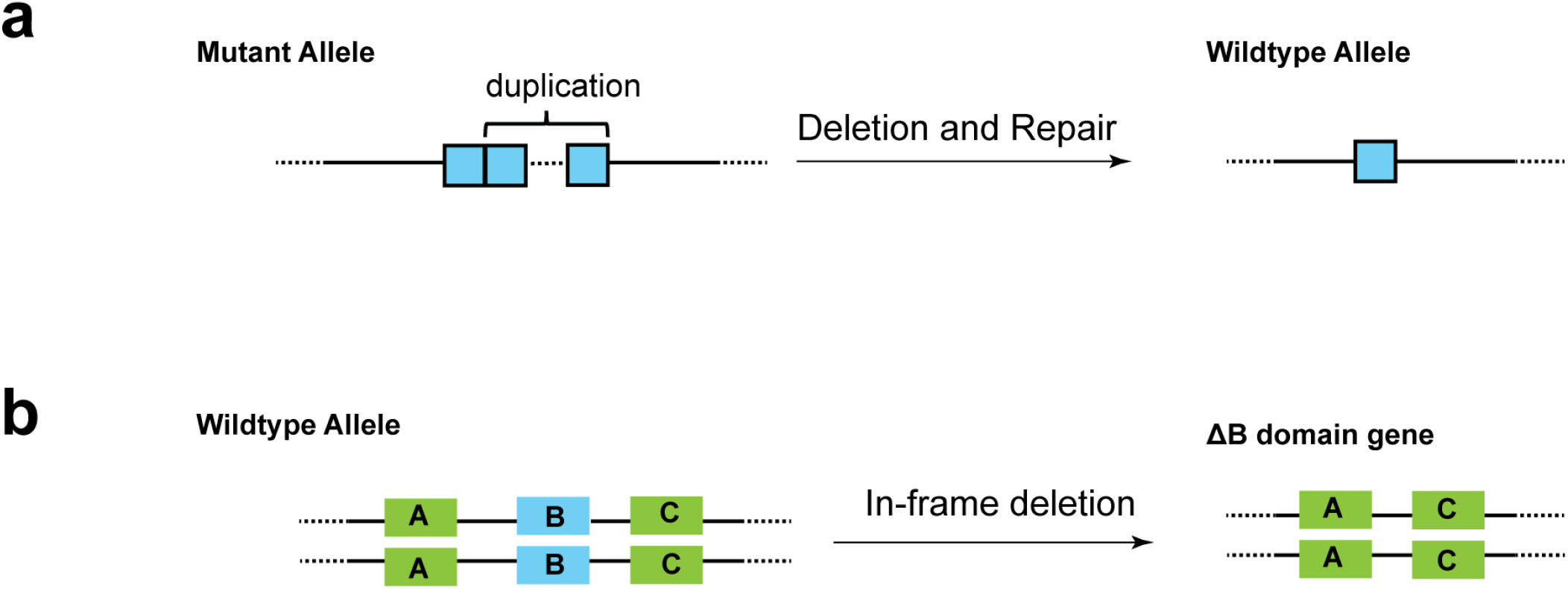
**a,** PEDAR could be used to delete a duplicated sequence to correct pathogenic mutations. **b,** PEDAR could also be used to study the functional domain of a protein by programming an in-frame deletion.

**Supplementary Fig. 8.**
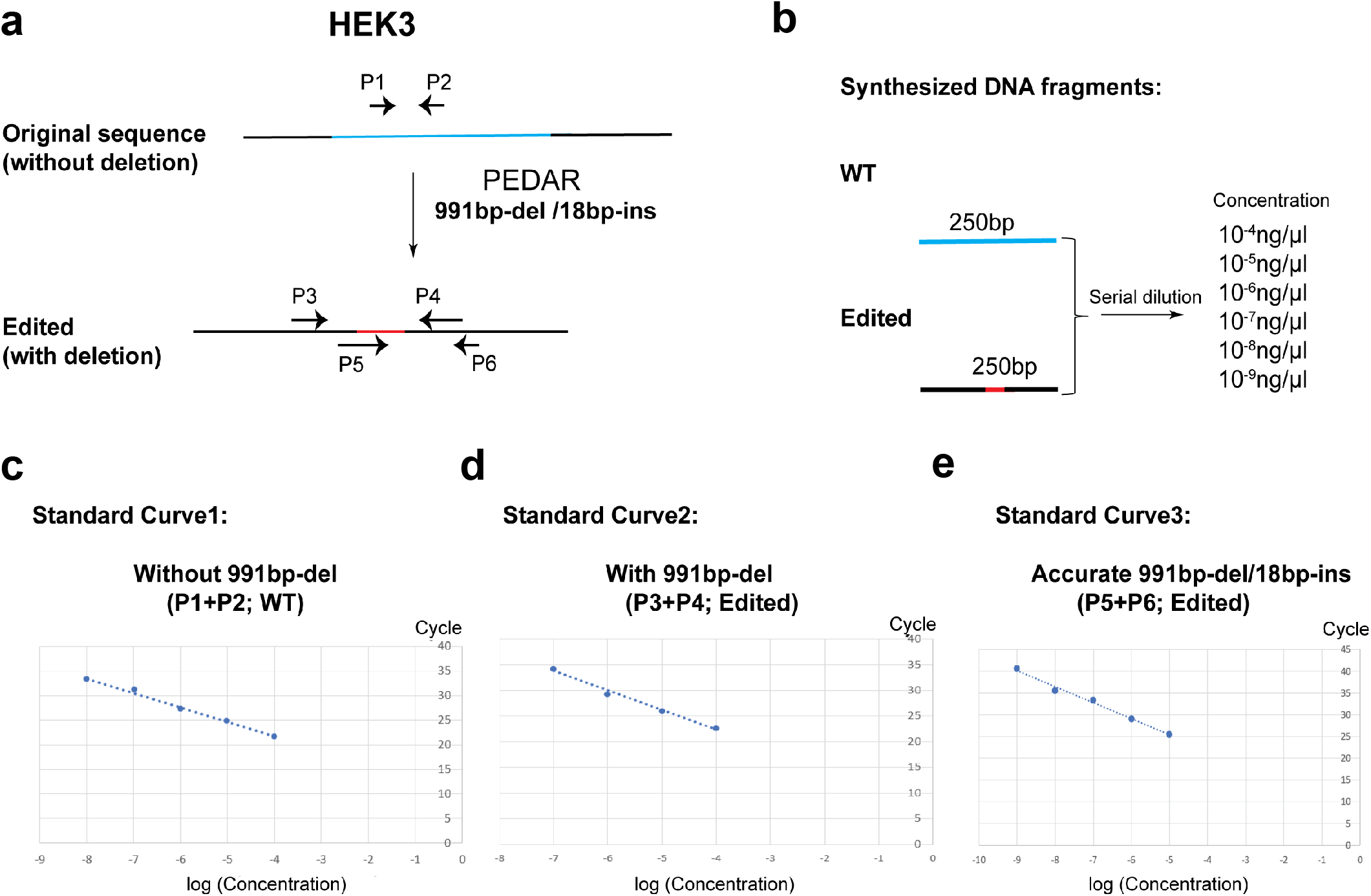
**a**, Design three pairs of qPCR primers to amplify the target site at the *HEK3* locus. b, Design two 250-bp DNA fragments (denoted as “WT” and “Edited”) of the same sequence with unedited or accurately edited target site. The two fragments were then serially diluted, serving as templates for standard curve. **c, d, e**, Three standard curves are generated by quantitative PCR using indicated paired primers and templates. Curves reflect the correlation between qPCR cycle number and the concentration of DNA without the 991-bp deletion (c), with the 991-bp deletion (d), or with the accurate 991-bp deletion/18-bp insertion (e).

